# Functional redundancy between penicillin-binding proteins during asymmetric cell division in *Clostridioides difficile*

**DOI:** 10.1101/2024.09.26.615255

**Authors:** Shailab Shrestha, Jules M. Dressler, Gregory A. Harrison, Morgan E. McNellis, Aimee Shen

## Abstract

Peptidoglycan synthesis is an essential driver of bacterial growth and division. The final steps of this crucial process involve the activity of the SEDS family glycosyltransferases that polymerize glycan strands and the class B penicillin-binding protein (bPBP) transpeptidases that cross-link them. While many bacteria encode multiple bPBPs to perform specialized roles during specific cellular processes, some bPBPs can play redundant roles that are important for resistance against certain cell wall stresses. Our understanding of these compensatory mechanisms, however, remains incomplete. Endospore-forming bacteria typically encode multiple bPBPs that drive morphological changes required for sporulation. The sporulation-specific bPBP, SpoVD, is important for synthesizing the asymmetric division septum and spore cortex peptidoglycan during sporulation in the pathogen *Clostridioides difficile*. Although SpoVD catalytic activity is essential for cortex synthesis, we show that it is unexpectedly dispensable for SpoVD to mediate asymmetric division. The dispensability of SpoVD’s catalytic activity requires the presence of its SEDS partner, SpoVE, and is facilitated by another sporulation-induced bPBP, PBP3. Our data further suggest that PBP3 interacts with components of the asymmetric division machinery, including SpoVD. These findings suggest a possible mechanism by which bPBPs can be functionally redundant in diverse bacteria and facilitate antibiotic resistance.

**Importance:** Peptidoglycan synthesis requires the transpeptidase activity of penicillin-binding proteins (PBPs), which can have specialized functions during cellular growth, division, and differentiation. However, many bacteria produce PBPs with overlapping functions, and this functional redundancy can increase antibiotic resistance. The spore-forming pathogen, *Clostridioides difficile*, encodes four PBPs: two are essential for growth and division and another, SpoVD, is essential for spore formation. Here, we show that the transpeptidase activity of SpoVD is dispensable during the first morphological step of sporulation, asymmetric division, because a sporulation-induced PBP, PBP3, partially substitutes for SpoVD’s function during this early stage. We find that SpoVD and PBP3 interact during sporulation, suggesting a mechanism by which bPBPs can confer functional redundancy and potentially contribute to antibiotic resistance.

## Introduction

Peptidoglycan (PG) synthesis is an essential driver of the morphological changes required for bacterial growth and division. This crucial process is driven by the glycosyltransfer reactions that polymerize the glycan strands and the transpeptidation reactions that crosslink the peptide sidechains between the strands (Egan et al., 2020; Rohs & Bernhardt, 2021). These enzymatic reactions require the activities of high-molecular-weight (HMW) Penicillin-binding proteins (PBPs), which are divided into two classes based on their catalytic ability: class A PBPs (aPBP) are bifunctional enzymes capable of both glycosyltransferase and transpeptidase activities, while class B PBPs (bPBPs) are monofunctional transpeptidases (Goffin & Ghuysen, 1998; Sauvage et al., 2008). Since crosslinking of PG is essential in almost all bacteria, inhibiting PBP transpeptidase activity is typically lethal to bacterial cells. Consequently, beta-lactam antibiotics such as penicillin, which inhibit PBPs by covalently bonding to the catalytic serine residue in their transpeptidase domain, are some of the most successful and widely used antibiotics (Zapun et al., 2008). As such, identifying factors that confer resistance to beta-lactam antibiotics and determining their mechanism of action has been an area of significant interest.

One mechanism through which bacteria achieve resistance against beta-lactam antibiotics is by inducing the production of PBPs with lower binding affinities for specific beta-lactams.

Most bacteria encode multiple PBPs that are specialized for specific cellular processes; some PBPs are essential, while others can be functionally redundant (Goffin & Ghuysen, 1998; Sauvage et al., 2008). Essential PBPs typically function as core components of highly conserved multiprotein complexes that drive cell wall synthesis during growth and division. The divisome is the essential complex that drives septal PG synthesis through the activities of a bPBP transpeptidase that is partnered with a cognate glycosyltransferase of the shape, elongation, division, and sporulation (SEDS) protein family (Cameron & Margolin, 2024; Rohs & Bernhardt, 2021; Taguchi et al., 2019). The elongasome is a multiprotein complex that drives cell elongation in rod-shaped bacteria through the action of a distinct SEDS-bPBP pair (Emami et al., 2017; Meeske et al., 2015; Rohs & Bernhardt, 2021; Sjodt et al., 2020). Notably, these SEDS-bPBP pairs are highly specific because the bPBP acts as a selective allosteric activator of its cognate SEDS family glycosyltransferase (Shlosman et al., 2023; Sjodt et al., 2020) (**Figure 1a**).

**Figure 1.**
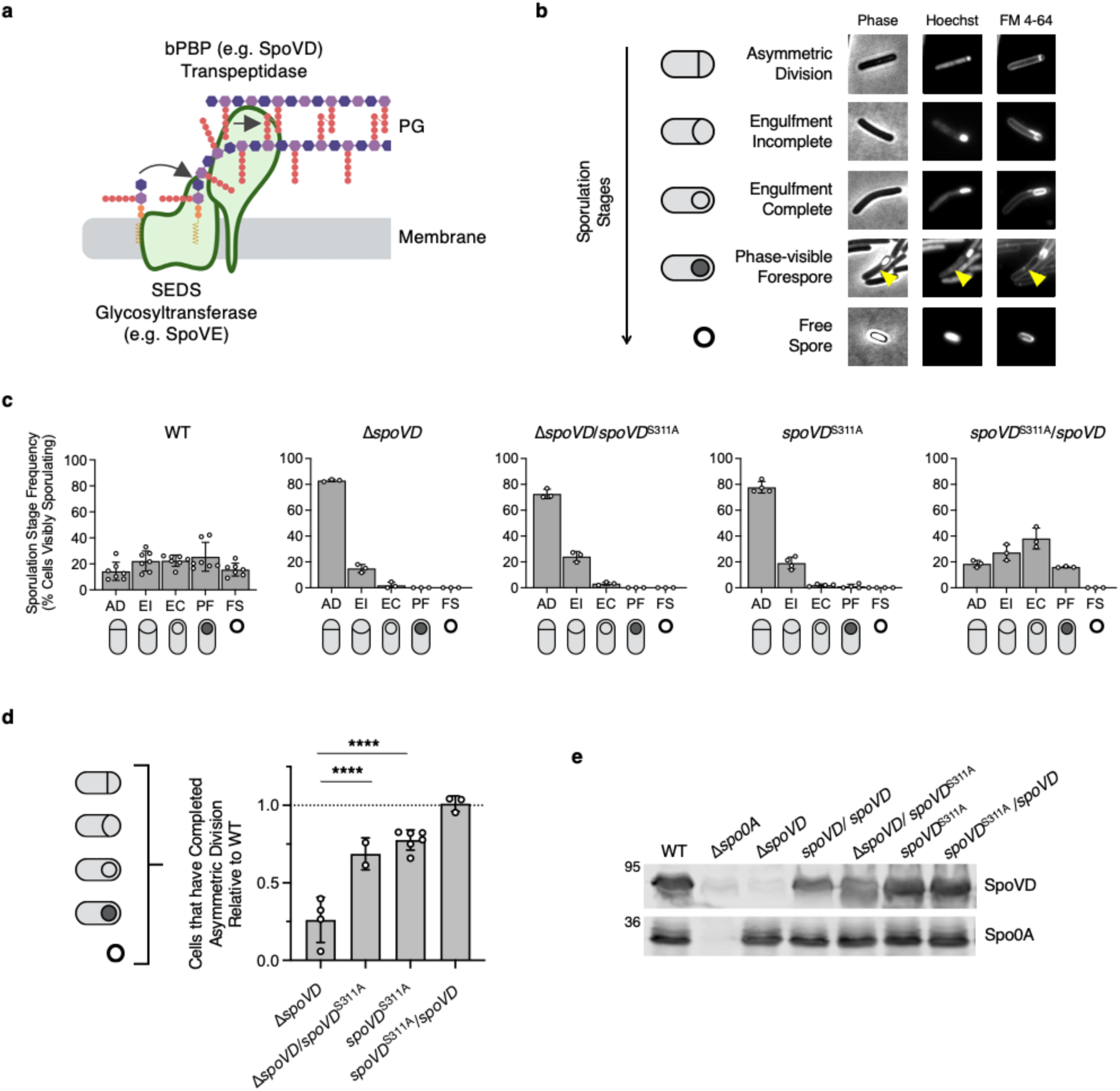
SpoVD catalytic activity is partially dispensable for its function during asymmetric division. **a** Schematic of a SEDS-bPBP peptidoglycan synthase complex. The SEDS glycosyltransferase polymerizes nascent glycan strands from lipid-linked PG precursors in the cytoplasmic membrane, while the class B PBP (bPBP) crosslinks the stem peptides between the growing strands. **b** Cytological profile of individual cells representing each of the five morphological stages of sporulation as indicated. Representative phase-contrast and fluorescence micrographs are WT cells sampled from sporulation-inducing 70:30 plates after 18 hours of growth. The nucleoid was stained using Hoechst, and the cell membrane was stained using FM4-64. Cells undergoing asymmetric division (AD) have a flat polar septum; cells undergoing engulfment (EI) have a curved polar septum; cells that have completed engulfment (EC) are indicated by bright-membrane staining around a fully engulfed forespore; Phase-visible forespores (PF) indicate forespores completing maturation visible as phase-dark or phase-bright forespores (yellow arrow) associated with the mother cell; mature free spores (FS) are observable as independent phase-bright particles. **c, d** Quantification of the cytological profiling of cells sampled from sporulation-inducing plates after 20-22 hours of growth. White circles indicate data from each replicate, bars indicate the average means, and error bars indicate standard deviation. >1,000 total cells and >100 visibly sporulating cells per sample from a minimum of three biological replicates. **c** shows the distribution of visibly sporulating cells among the indicated stages of sporulation. **d** shows the proportion of cells that complete and progress beyond asymmetric division, i.e., all visibly sporulating cells, as a percentage of the total cells profiled. Note that the data is normalized to WT (dotted line). **** p < 0.0001. Statistical significance was determined using a one-way ANOVA and Tukey’s test. **e** Western blot analyses of SpoVD levels in the indicated strains 14 hours after growth on sporulation-inducing plates. The anti-Spo0A antibody was used as a proxy for measuring sporulation induction.

Endospore-forming bacteria typically encode an additional SEDS-bPBP complex that is responsible for driving the morphological changes required for spore formation (Galperin et al., 2012, 2022; Shrestha et al., 2023; Tan & Ramamurthi, 2014). Sporulation begins with the formation of a polar division septum close to one cell pole in a process called asymmetric division. In *Bacillus subtilis*, asymmetric division is driven by the same SEDS-bPBP pair that mediates cell division during vegetative growth (Muchová et al., 2020; Real et al., 2008). In contrast, we recently showed that the spore-forming pathogen *Clostridioides difficile* lacks a canonical division-associated SEDS-bPBP pair for driving septal PG synthesis and instead uses an aPBP as the major PG synthase during vegetative cell division (Shrestha et al., 2023). In further contrast with *B. subtilis,* the sporulation-specific SEDS-bPBP pair SpoVE-SpoVD is an important driver of septal PG synthesis during asymmetric division in *C. difficile*. Although the role of SpoVE-SpoVD function during asymmetric division may be restricted to *C. difficile* and other clostridial organisms, genes encoding SpoVE and SpoVD can be found in almost all spore formers (Galperin et al., 2012, 2022). In both *B. subtilis* and *C. difficile*, SpoVE and SpoVD are essential for synthesizing the spore cortex, a thick layer of modified PG that surrounds and protects the spore core (Henriques et al., 1992; Yanouri et al., 1993; Daniel et al., 1994; Shrestha et al., 2023; Alabdali et al., 2021; Srikhanta et al., 2019). Thus, SpoVE and SpoVD are important for synthesizing PG during two distinct stages of spore formation in *C. difficile*.

While a previous study showed that SpoVD’s catalytic activity is essential for spore formation in *C. difficile* (Alabdali et al., 2021), in this study, we surprisingly observe that SpoVD catalytic activity is largely dispensable for mediating asymmetric division despite being essential for synthesizing the cortex layer. Prior analyses of a catalytic mutant of a divisome-associated bPBP, PBP2b*_Bs_*, in *B. subtilis* provide a possible mechanism for explaining this observation. In *B. subtilis*, the catalytic activity of PBP2b*_Bs_* is dispensable because a second bPBP, PBP3*_Bs_*, can supply the transpeptidase activity during septal PG synthesis (Sassine et al., 2017). Since the gene encoding PBP2b*_Bs_* is essential, presumably because the PBP2b*_Bs_* protein is required to allosterically activate the glycosyltransferase activity of its SEDS binding partner, FtsW, these observations highlight that PBP3*_Bs_* cannot complement all the roles fulfilled by the catalytically inactive PBP2b*_Bs_*. Notably, PBP3*_Bs_* is also important for resistance against certain beta-lactams, since it has lower affinities for them compared to PBP2b*_Bs_* (Sassine et al., 2017). While these observations highlight the importance of catalytic redundancies between PBPs, the molecular mechanisms behind this phenomenon remain unclear.

Here, we explore the role of SpoVD’s catalytic activity during asymmetric division in *C. difficile*. Our findings suggest that the ability of a catalytically inactive SpoVD to support septal PG synthesis during asymmetric division requires the presence of its SEDS partner SpoVE and is facilitated in part by another sporulation-specific bPBP, PBP3. Furthermore, we provide evidence suggesting that PBP3 interacts with components of the asymmetric division machinery, including SpoVD. These findings suggest a possible mechanism for how bPBPs in diverse bacteria can be functionally redundant and promote antibiotic resistance.

## Results

### A SpoVD catalytic mutant is defective in cortex synthesis but not in asymmetric division

Given that the catalytic activity of the division-specific bPBP in *B. subtilis* is not essential for its role in cell division (Sassine et al., 2017), we speculated that the catalytic activity of SpoVD might be similarly dispensable during asymmetric division in *C. difficile*. SpoVD retains the SXXK active-site motif that is highly conserved among bPBPs. To abolish the catalytic activity of SpoVD, we introduced an alanine substitution at the nucleophilic serine residue (SpoVD^S311A^). Consistent with a prior study (Alabdali et al., 2021), *C. difficile* strains with a catalytically inactive SpoVD failed to produce heat-resistant spores (**Figure S1**). Importantly, this phenotype was observed when *spoVD*^S311A^ was expressed from an ectopic chromosomal locus (Δ*spoVD*/*spoVD*^S311A^) or the native locus (*spoVD*^S311A^) (**Figure S1**).

To further define the role of SpoVD’s catalytic activity during spore formation, we evaluated the ability of the SpoVD catalytic mutant strains to progress through the different morphological stages of sporulation by cytologically profiling sporulating cells (**Figures 1b** and **S2**) (Nonejuie et al., 2013; Pogliano et al., 1999). Similar to the *spoVD* deletion strain, *spoVD* catalytic mutant strains failed to produce phase-visible spores, indicating that they have defects in cortex synthesis (**Figure 1c**). Furthermore, the distribution of sporulating cells among different stages was similar between the strains, with most cells being stalled at the asymmetric division stage and relatively few cells completing engulfment (**Figure 1c**). To specifically quantify the effects on asymmetric division, we calculated the proportion of cells that showed morphological signs of sporulation in the cytological profiling, i.e., cells that progressed beyond asymmetric division, as a percentage of all cells. These analyses revealed that, relative to WT, Δ*spoVD*/*spoVD*^S311A^ and *spoVD*^S311A^ cells complete asymmetric division at significantly higher frequencies compared to Δ*spoVD* cells, with Δ*spoVD* cells completing asymmetric division at ∼30% the levels of WT and Δ*spoVD*/*spoVD*^S311A^ and *spoVD*^S311A^ cells completing asymmetric division at ∼70% the levels of WT (**Figure 1d**). Since we previously showed that the decreased frequency of visibly sporulating Δ*spoVD* cells reflects a defect in asymmetric division rather than sporulation initiation (Shrestha et al., 2023), these results suggest that SpoVD function during asymmetric division is only partially dependent on its catalytic activity. Importantly, the levels of SpoVD present in these strains upon sporulation induction were found to be similar to wild-type (WT) by western blot analysis (**Figure 1e**).

Surprisingly, while we were able to complement the asymmetric division defects of the catalytic mutant strains by expressing a wild-type copy of *spoVD* from an ectopic locus (**Figure 1c, 1d**), this strain was unable to form mature phase-bright spores (**Figure S1, 1c**). This suggests that the catalytically inactive SpoVD has a dominant negative phenotype specifically for the defect in cortex synthesis.

### The SpoVD catalytic mutant requires its SEDS partner, SpoVE

Since previous studies suggest that bPBPs are needed to allosterically activate the glycosyltransferase activity of their cognate SEDS family glycosyltransferases (Shlosman et al., 2023; Sjodt et al., 2020; Taguchi et al., 2019), we tested if the ability of SpoVD^S311A^ to support asymmetric division requires its SEDS partner SpoVE. To this end, we created a *spoVD*^S311A^Δ*spoVE* strain by introducing the catalytic mutant variant of *spoVD* into the native locus of a Δ*spoVD*Δ*spoVE* strain. As expected, this strain failed to form heat-resistant spores (**Figure 2**). Cytological profiling of sporulating cells revealed that *spoVD*^S311A^Δ*spoVE* cells complete and progress beyond asymmetric division at a significantly lower frequency (∼25%) than WT or *spoVD*^S311A^ cells (100% and ∼75%, respectively). Thus, the phenotype of *spoVD*^S311A^Δ*spoVE* cells is similar to that of Δ*spoVD* and Δ*spoVE* cells (**Figures 1c, d & 2a, b**). Since SpoVD levels are unaffected by loss of SpoVE (**Figure 2c**), consistent with our prior results (Shrestha et al., 2023), we conclude that the function of SpoVD^S311A^ during asymmetric division requires SpoVE.

**Figure 2.**
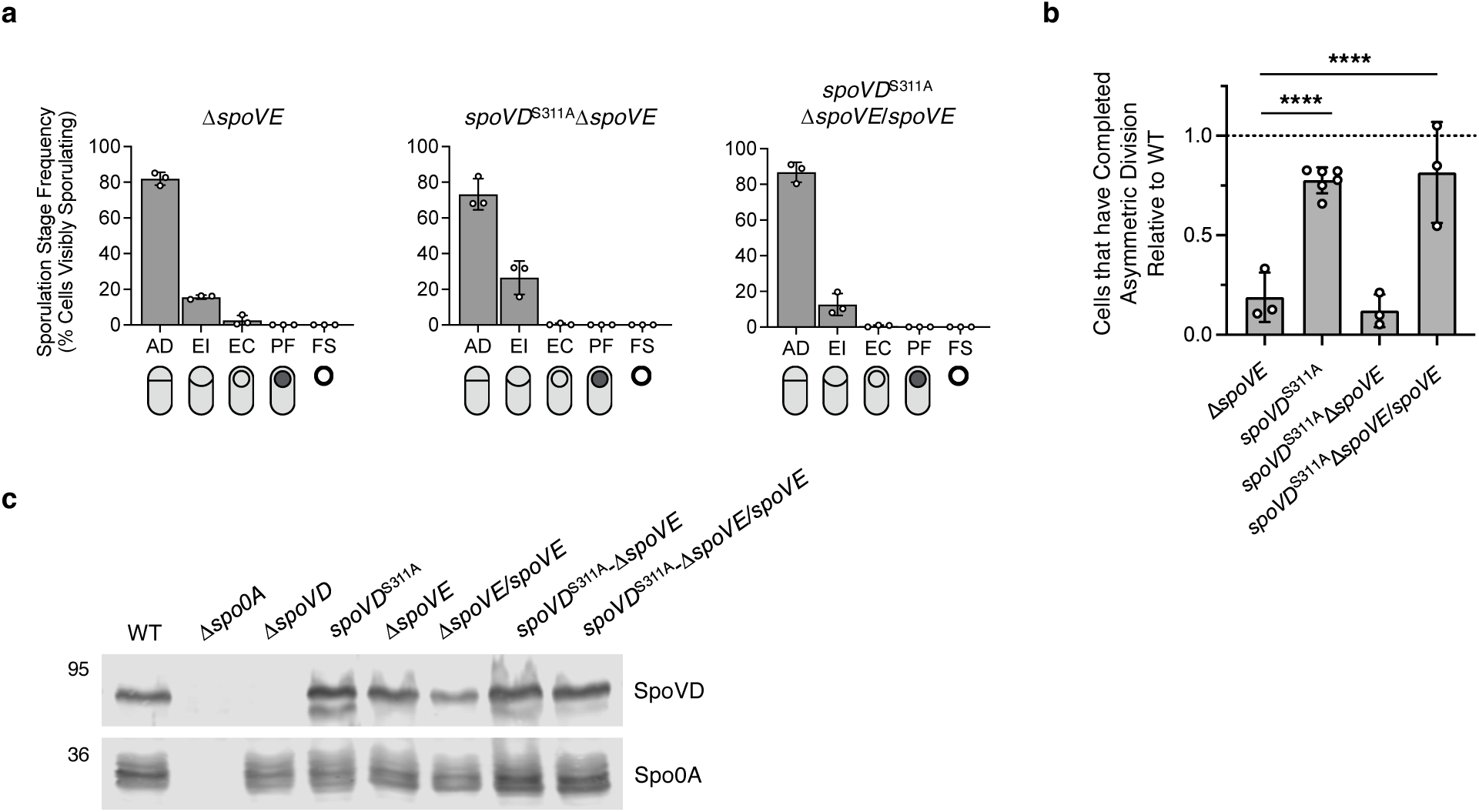
The catalytically inactive SpoVD requires its SEDS partner, SpoVE, to facilitate asymmetric division. **a, b** Quantification of the cytological profiling of cells sampled from sporulation-inducing plates after 20-22 hours of growth. White circles indicate data from each replicate, bars indicate average means, and error bars indicate standard deviation. >1,000 total cells and >100 visibly sporulating cells per sample from a minimum of three biological replicates. **a** shows the distribution of visibly sporulating cells among the indicated stages of sporulation. See Figure 1 for the distribution of WT cells. **b** shows the proportion of cells that complete and progress beyond asymmetric division, i.e., all visibly sporulating cells, as a percentage of the total cells profiled. Note that the data is normalized to WT (dotted line) and that the *spoVD^S311A^* data derives from Figure 1. *** p < 0.001, **** p < 0.0001. Statistical significance was determined using a one-way ANOVA and Tukey’s test. **c** Western blot analyses of SpoVD levels in the indicated strains 14 hours after growth on sporulation-inducing plates. The anti-Spo0A antibody was used as a proxy for measuring sporulation induction.

### An additional non-essential bPBP, PBP3, is involved in sporulation

Next, we wondered if *C. difficile* encodes an additional sporulation-specific bPBP that can cross- link septal PG synthesis in the absence of SpoVD catalytic activity similar to the functional redundancy observed between division-specific bPBPs in *B. subtilis* (Sassine et al., 2017). Most *C. difficile* strains encode three bPBPs (Isidro et al., 2017, 2018): the sole essential bPBP, PBP2, primarily functions during cell elongation, while the two other bPBPs, SpoVD and PBP3, are dispensable for vegetative growth (Shrestha et al., 2023). PBP3 is a close homolog of SpoVD containing a similar domain composition apart from the C-terminal PASTA (PBP And Serine/Threonine kinase Associated) domain carried by SpoVD (**Figure 3a**). Both PBP3 and SpoVD have an N-terminal transmembrane section followed by PBP dimerization and transpeptidase domains.

**Figure 3.**
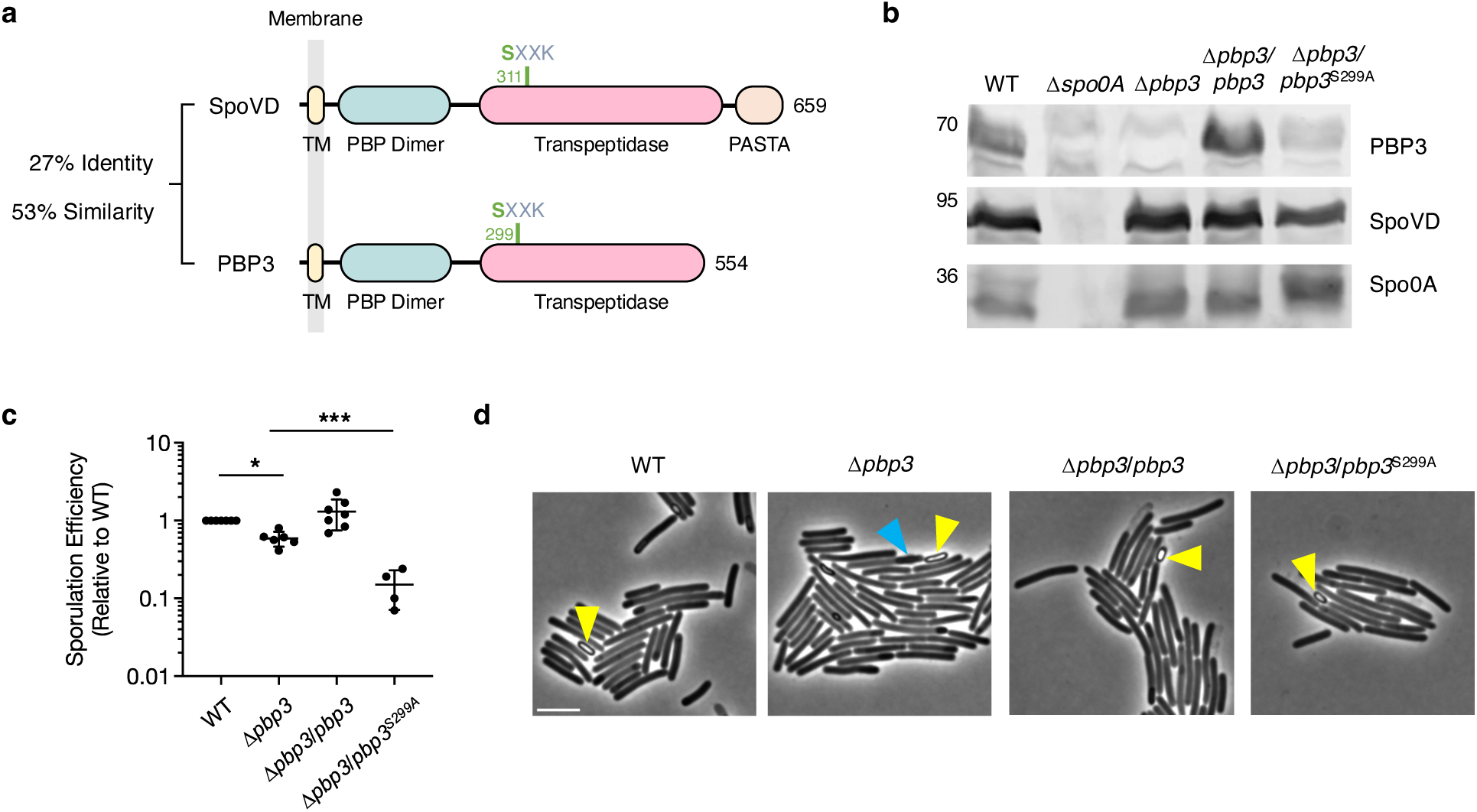
PBP3 is a sporulation-specific bPBP that is involved in spore formation. **a** Protein schematic comparing SpoVD and PBP3. Functional domains and catalytic sites were predicted using HMMER. TM, transmembrane domain; PASTA, PBP, and serine/threonine kinase-associated domain. The catalytic serine residues are shown in green. **b** Western blot showing the levels of the SpoVD, PBP3, and Spo0A in cells sampled from sporulation-inducing plates after ∼14 hours of growth. SpoVD and PBP3 are not detected in the Δ*spo0A* strain, which cannot initiate sporulation. **c** Efficiency of heat-resistant spore formation (sporulation efficiency) of the *pbp3* mutant and complemented strains relative to WT. Means with standard deviation are indicated. Cells were collected from sporulation-inducing 70:30 plates ∼20-22 hours after inoculation. Data from a minimum of four biological replicates. * p < 0.05, *** p < 0.001. Statistical significance was determined using a one-way ANOVA and Tukey’s test. **d** Representative phase-contrast micrographs of WT, *pbp3* mutant, and complemented cells collected from sporulation-inducing 70:30 plates after ∼20 hours of growth. Examples of phase-bright spores are indicated by yellow arrows. The blue arrow highlights an elongated forespore in the Δ*pbp3* mutant. Scale bar, 5 μm.

We considered PBP3 to be a likely candidate for providing functional redundancy to SpoVD during asymmetric division for several reasons. First, previous studies suggest that, similar to *spoVD* expression, the expression of *pbp3* (locus *cd630_12290* in the strain used in this study) is upregulated at the onset of sporulation (Fimlaid et al., 2013; Saujet et al., 2013). Second, we previously showed that individual deletions of *spoVD* or *pbp3* do not affect the growth rate of vegetative cells, suggesting that SpoVD and PBP3 play sporulation-specific roles (Shrestha et al., 2023). This agrees with our western blot analysis showing that SpoVD and PBP3 are produced under sporulation-inducing conditions in a manner dependent on the presence of the master transcriptional regulator of sporulation, Spo0A (**Figure 3b**). Taken together with prior transcriptomic studies (Fimlaid et al., 2013; Saujet et al., 2013), which similarly show Spo0A- dependent induction of *pbp3* and *spoVD* transcription, these observations demonstrate that both proteins are produced prior to asymmetric division.

Further characterization of the Δ*pbp3* strain revealed that it forms heat-resistant spores ∼2-fold less efficiently than WT, a modest defect that can be complemented by expressing *pbp3* from an ectopic chromosomal locus (**Figure 3c**). This result is consistent with a prior transposon mutagenesis screen, which identified *pbp3* to be important for sporulation (Dembek et al., 2015). Phase-contrast microscopy of sporulating cells revealed that the Δ*pbp3* mutant forms phase- bright forespores and free spores (**Figure 3d** and **S3**), indicating that it can assemble the spore cortex unlike a Δ*spoVD* mutant. Consistent with this conclusion, transmission electron microscopy analyses detected the cortex layer in Δ*pbp3* forespores (**Figure S4**). Interestingly, these analyses also revealed that the Δ*pbp3* mutant frequently makes forespores that are elongated and slightly misshapen, indicating that PBP3 affects spore formation downstream of asymmetric division (**Figure S4**). The slight morphological defects may explain the reduction in heat-resistant spore formation relative to WT cells. Notably, while complementation of the Δ*pbp3* mutant with WT *pbp3* restored sporulation efficiency to WT levels, a strain complemented with a construct encoding a catalytic mutant (*pbp3*^S299A^) exhibited a decrease in heat-resistant spore formation, indicating that PBP3’s catalytic activity is important for proper spore formation (**Figure 3d**). Taken together, our results indicate that PBP3 is a sporulation-induced factor that is involved in, but not essential for, the formation of mature spores in *C. difficile*.

### PBP3 promotes asymmetric division in the absence of SpoVD catalytic activity during asymmetric division

To test if PBP3 provides redundancy to SpoVD catalytic activity during asymmetric division, we analyzed the ability of Δ*pbp3* cells to complete asymmetric division in the context of WT and catalytically inactive SpoVD variants. Cells lacking PBP3 completed asymmetric division at WT levels in the presence of WT SpoVD (**Figure 4b**). The distribution of sporulating Δ*pbp3* cells among the different sporulation stages was also similar to WT (**Figure 4a**). Importantly, in the *spoVD*^S311A^ background, cells lacking PBP3 completed asymmetric division at a lower rate when compared to WT or *spoVD*^S311A^ cells (**Figure 4b**). However, the proportion of *spoVD*^S311A^Δ*pbp3* cells that complete asymmetric division was higher compared to Δ*spoVD* cells (**Figure 4b**), suggesting that PBP3 partially accounts for the ability of the catalytically dead SpoVD to support asymmetric division. To establish if this partial redundancy requires the transpeptidase activity of PBP3, we introduced *pbp3*^S299A^, which encodes a catalytically inactive PBP3 in an ectopic chromosomal locus of a *spoVD*^S311A^ Δ*pbp3* strain. Cytological profiling of the *spoVD*^S311A^ Δ*pbp3*/*pbp3*^S299A^ strain revealed that these cells complete asymmetric division at levels similar to WT (**Figure 4**). Hence, our data suggest that PBP3 enhances asymmetric division in the absence of SpoVD catalytic activity during asymmetric division through a mechanism that is independent of PBP3’s transpeptidase activity.

**Figure 4.**
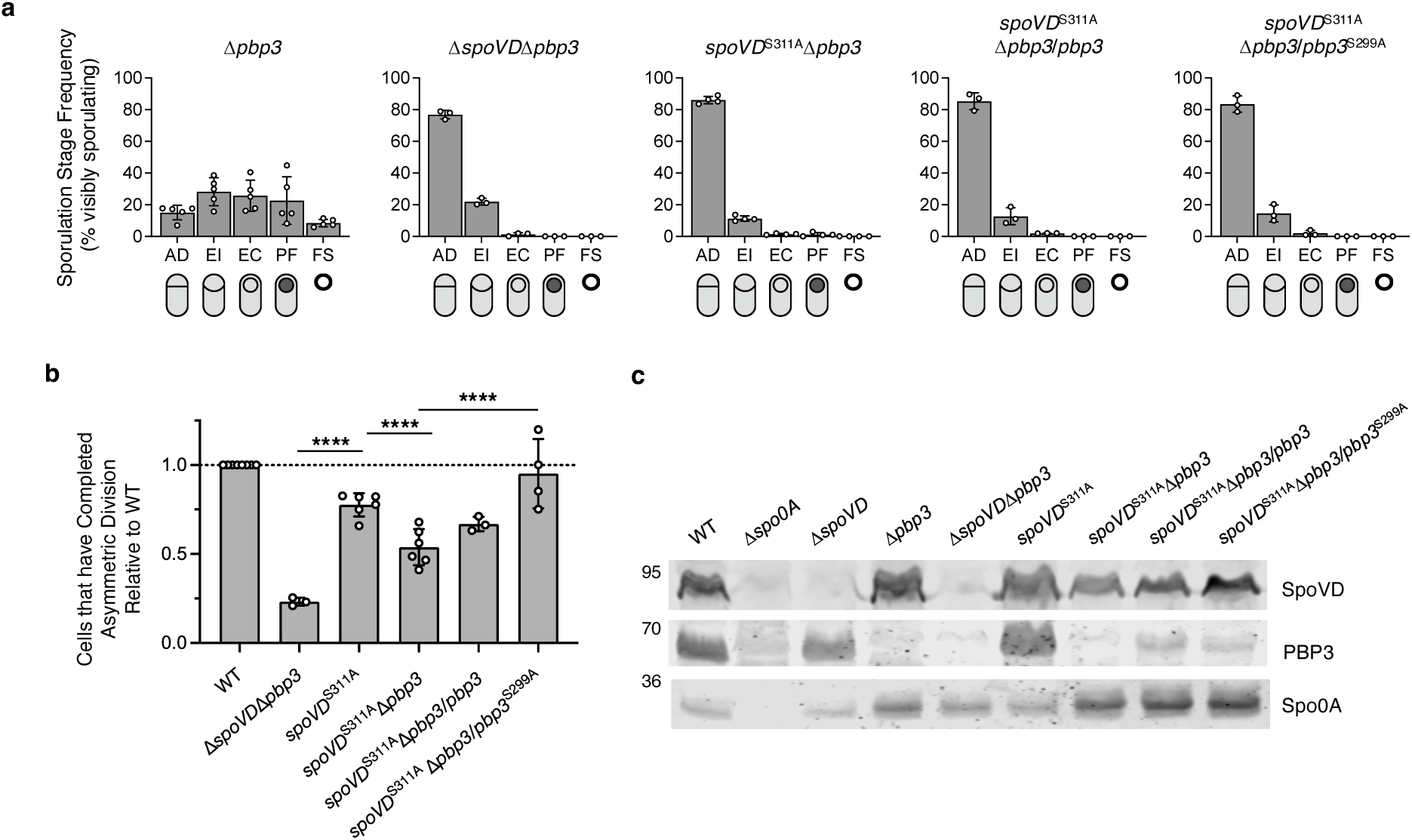
PBP3 partially compensates for loss of SpoVD catalytic activity during asymmetric division. **a, b** Quantification of the cytological profiling of cells sampled from sporulation-inducing plates after 20-22 hours of growth. White circles indicate data from each replicate, bars indicate average means, and error bars indicate standard deviation. >1,000 total cells and >100 visibly sporulating cells per sample. Data from a minimum of three independent experiments. **** p < 0.0001. Statistical significance was determined using a one-way ANOVA and Tukey’s test. **a** shows the distribution of visibly sporulating cells among the indicated stages of sporulation. See Figure 1b for the distribution of WT cells. **b** shows the proportion of cells that complete and progress beyond asymmetric division, i.e., all visibly sporulating cells, as a percentage of the total cells profiled. Note that the data is normalized to WT (dotted line) and that the *spoVD^S311A^* data derives from Figure 1. **c** Western blot showing the levels of the SpoVD, PBP3, and Spo0A in cells sampled from sporulation-inducing plates after ∼14 hours of growth. SpoVD and PBP3 are not detected in the Δ*spo0A* strain, which cannot initiate sporulation.

### PBP3 may interact with PG-synthesizing enzymes and components of the polar divisome

Our previous study suggests that SpoVD likely functions as part of the polar divisome to synthesize septal PG during asymmetric division (Shrestha et al., 2023). The partial redundancy between PBP3 and SpoVD catalytic activities during asymmetric division observed here suggests that PBP3 might be recruited to the asymmetric division machinery during sporulation. To test whether PBP3 interacts with components of the polar divisome, we conducted bacterial two-hybrid assays to probe pairwise interactions between PBP3 and known components of this complex. In addition to SpoVD and SpoVE, we explored interactions with three additional proteins, FtsL, FtsQ, and FtsB (also known as FtsL, DivIB, and DivIC in some Firmicutes), which form a highly-conserved divisome sub-complex that regulates PG synthase activity during cell division in other bacteria (Levin & Losick, 1994; Daniel et al., 1998; Katis & Wake, 1999; Tsang & Bernhardt, 2015; Marmont & Bernhardt, 2020; Shrestha et al., 2023). Our data suggests that PBP3 can interact with various components of the polar divisome, including SpoVD, SpoVE, and FtsQ (**Figure 5a**). As controls, we also probed interactions between PBP3 and all other PBP and SEDS-family proteins encoded by *C. difficile*. Surprisingly, our analysis indicates that PBP3 also interacts with PBP2 and PBP1 (**Figure 5a**), which primarily function during vegetative growth but likely also play important roles during spore formation. Although surprising, PBP3 may be capable of forming a complex with all three of the other PBPs in *C. difficile*.

**Fig. 5.**
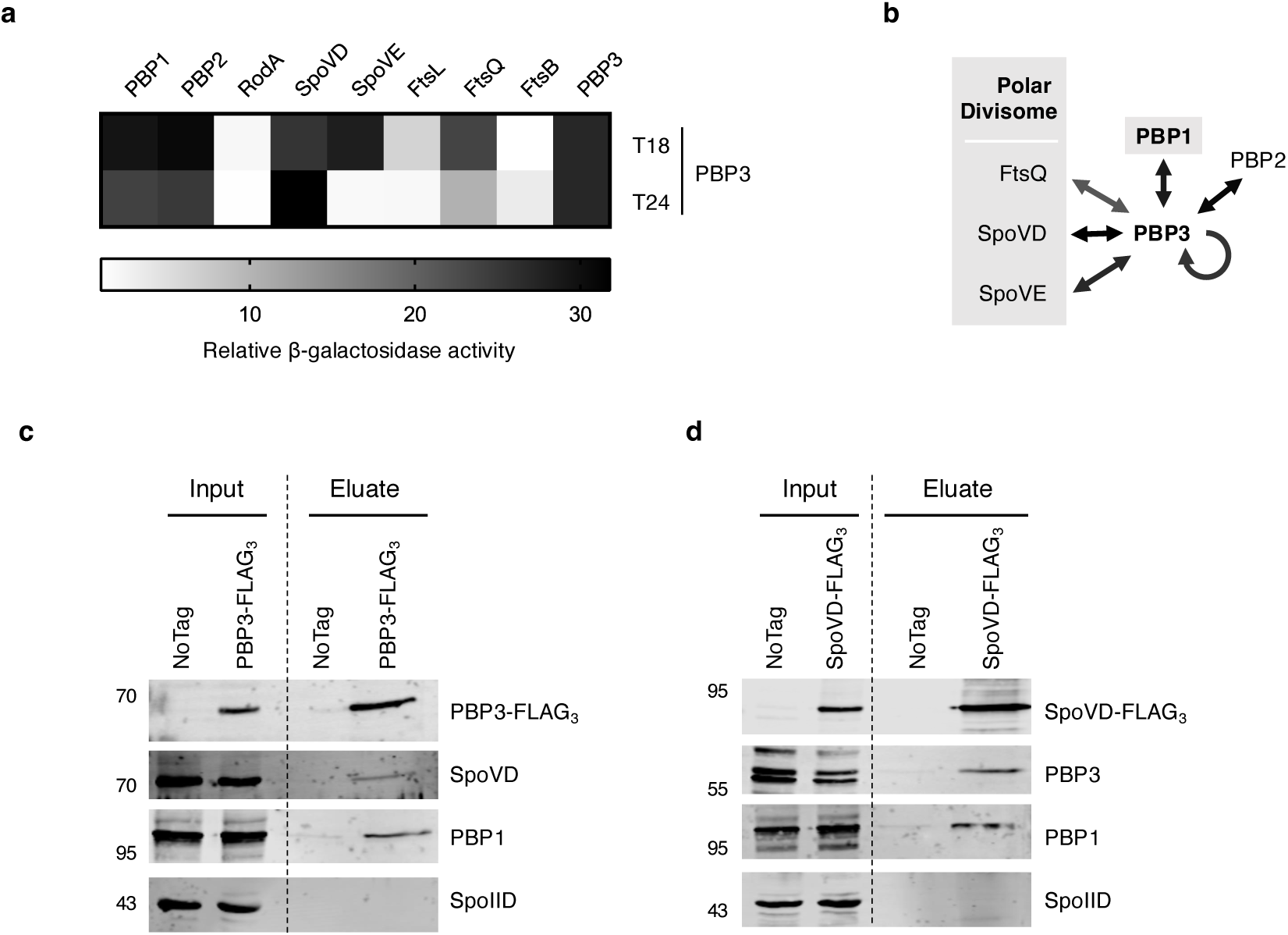
PBP3 interacts with multiple components of the polar divisome. **a** Bacterial two-hybrid analysis of interactions between PBP3 and other PG synthases or components of the polar divisome. The β-galactosidase activity was normalized to the negative control. N-terminal T18 or T24 fusion to PBP3 was paired with reciprocal N-terminal fusions to the indicated proteins. Data from three technical replicates. **b** The schematic shows interactions detected in the bacterial two-hybrid analyses. Components of the predicted polar divisome are indicated. PBP1 may also be a part of the polar divisome based on the co-immunoprecipitation analyses using SpoVD-FLAG3 as bait. **c, d** Co-immunoprecipitations performed on cells sampled from sporulation-inducing plates after 12 hours of growth. **c** PBP3-FLAG3 was used as bait in the Δ*pbp3/pbp3-FLAG3* strain background; **d** SpoVD-FLAG3 was used as bait in the Δ*spoVD/spoVD-FLAG3* strain background. Δ*pbp3/pbp3* and Δ*spoVD/spoVD* strains were used as negative controls (No Tag). The presence of SpoVD, PBP3, and PBP1 in the pull-downs was probed using antibodies against the indicated proteins and western blotting. The FLAG-tagged proteins were detected using an anti-FLAG antibody. SpoIID was used as a control protein because it is also a PG-associated transmembrane protein localized to the forespore membrane, but it is not predicted to be a part of the polar divisome.

Since bacterial two-hybrid analyses between two membrane proteins can lead to false- positive results likely due to high levels of tagged proteins in the membrane non-specifically reconstituting the adenylate cyclase enzyme (Yahashiri et al., 2020), we further tested whether PBP3 and SpoVD interact during *C. difficile* sporulation when expressed from their native promoters using co-immunoprecipitation analyses. To this end, we generated Δ*pbp3* and Δ*spoVD* strains complemented with constructs encoding FLAG-tagged PBP3 and FLAG-tagged SpoVD, respectively. Cell lysates were prepared from cultures grown on sporulation media for 12 hours when asymmetric division peaks in sporulating cells (Oliveira et al., 2020), and the FLAG- tagged PBPs were pulled-down. Western blot analyses of the pull-downs revealed that SpoVD was specifically enriched in PBP3-FLAG_3_ pull-downs (**Figure 5c**), and PBP3 was enriched in SpoVD-FLAG_3_ pull-downs (**Figure 5d**). Interestingly, PBP1, which is required for mediating vegetative cell division but also likely contributes to asymmetric division (Shrestha et al., 2023), was also specifically enriched in the pull-downs, suggesting that PBP1 can interact with one or both SpoVD and PBP3, directly or indirectly. Importantly, a control membrane-bound sporulation protein, SpoIID (Ribis et al., 2018), which is a peptidoglycan hydrolase that forms part of the “engulfasome” (Kelly & Salgado, 2019), was not found in the pull-downs. Overall, our data indicate that PBP3 is recruited to the polar divisome complex through direct interactions with its components (**Figure 5b**).

### Endospore-forming bacteria typically encode multiple additional bPBPs compared to non- sporulating bacteria

Since *B. subtilis* also encodes multiple sporulation-specific class B PBPs, we wondered whether functional redundancies between sporulation-specific bPBPs might be more broadly conserved in endospore-forming bacteria. To this end, we compared the numbers of SEDS and bPBP enzymes encoded in the genomes of non-sporulating and sporulating Firmicutes, the sole bacterial phylum with endospore-forming members. Our analyses revealed that sporulating bacteria typically encode a higher number of bPBPs compared to non-sporulating bacteria (**Figure 6**). In sporulating organisms, the number of encoded bPBPs often exceeds the number of encoded SEDS (214/328; 65%), while only a small minority encode more SEDS proteins than bPBPs (20/328; 6%). Furthermore, a higher percentage of non-sporulating organisms encode equal numbers of SEDS and bPBP genes (80%). These observations are consistent with the prevalence of additional sporulation-specific PG synthases in spore formers and suggest that redundancy in bPBP activity is likely widespread in these organisms.

**Fig. 6.**
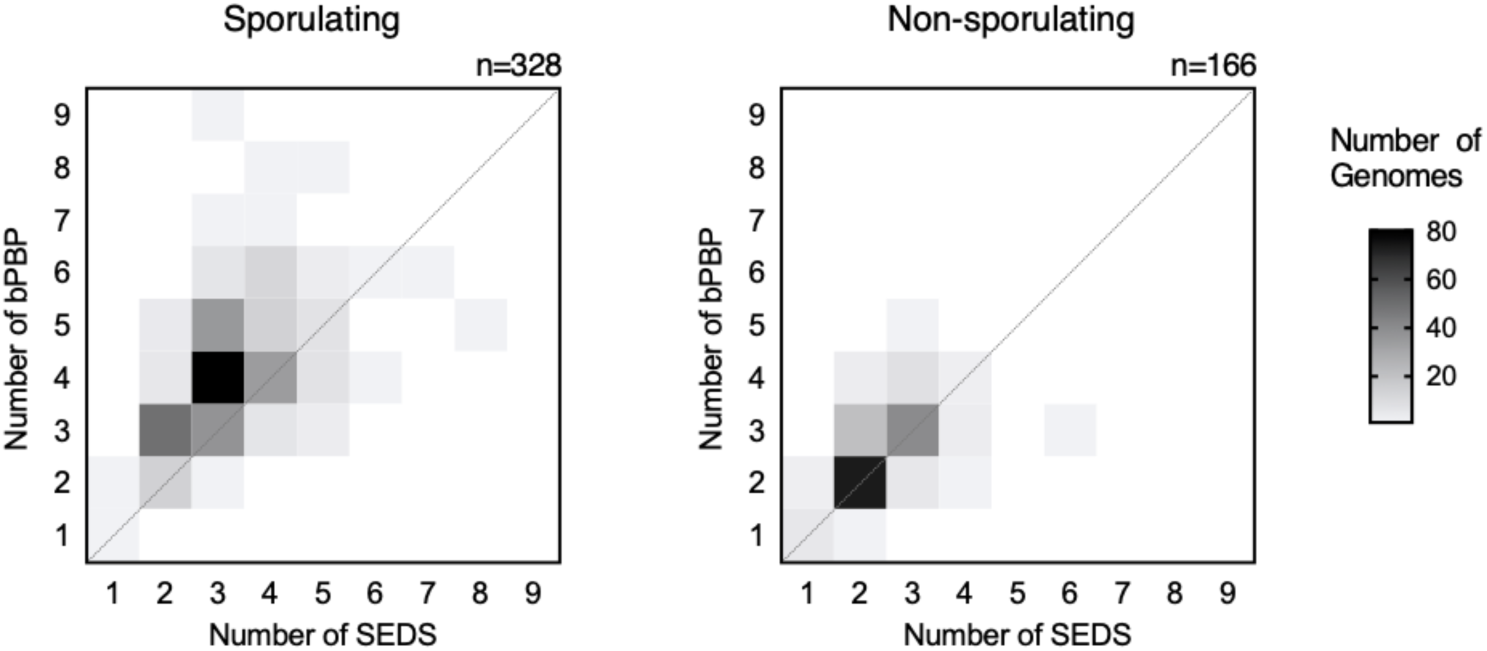
Prevalence of bPBP and SEDS enzymes in Firmicutes. Heatmaps showing the distribution of class B penicillin-binding protein (bPBP) and SEDS protein numbers encoded in the genomes of sporulating (n = 328) and non-sporulating (n = 166) Firmicutes organisms. Sporulation ability was inferred by the presence of broadly conserved sporulation-specific genes *spo0A* and *spoIIE* in the genome. The dataset comprises 494 diverse Firmicutes organisms, as reported in Shrestha *et al*., 2023.

## Discussion

While most bacteria encode multiple bPBPs that perform specialized roles during specific cellular processes, some bPBPs can play redundant roles that are important for resistance against environmental stresses. Our understanding of these compensatory mechanisms and their prevalence, however, remains incomplete. Here, we reveal that the catalytic activity of SpoVD in *C. difficile* is partially dispensable for its function during the synthesis of the polar septum (**Figure 1)**. Our data indicate that a previously uncharacterized sporulation-specific bPBP, PBP3, partially substitutes for loss of SpoVD’s catalytic activity during asymmetric division (**Figures 3, 4**), although PBP3’s catalytic activity is surprisingly dispensable for promoting asymmetric division in the absence of SpoVD’s catalytic activity (**Figure 4**). In contrast, PBP3’s catalytic activity is necessary for events downstream of asymmetric division to maximize functional spore formation (**Figure 3**). Finally, our bacterial two-hybrid and co-immunoprecipitation data suggest that PBP3 likely functions as a part of the polar divisome, which mediates asymmetric division (**Figure 5**).

Functional redundancies between PBPs affect the ability of bacteria to adapt to environmental stresses and resist antibiotics targeting PG transpeptidation. Redundancies between aPBPs have been described in *Escherichia coli* (Mueller et al., 2019), *B. subtilis,* and *Streptococcus pneumoniae* (Mitchell et al., 2023), where distinct aPBPs are specialized to respond to changes in environmental pH but can also play redundant roles in certain conditions. Similarly, niche specialization of different bPBPs was observed in *Salmonella enterica*, which encodes two division-specific and two elongation-specific bPBPs that are functionally specialized for extracellular or intracellular conditions (Castanheira et al., 2017, 2020).

Furthermore, as described above, functional redundancy in bPBP activities during cell division in *B. subtilis* lowers its sensitivity to certain beta-lactams (Sassine et al., 2017). Given that different bPBPs have distinct binding affinities to beta-lactams, bPBPs that are intrinsically resistant to certain beta-lactams can facilitate resistance by providing transpeptidation activity when essential bPBPs are inhibited. This is supported by studies in methicillin-resistant *Staphylococcus aureus* (MRSA), where strains that have acquired an additional low-affinity bPBP display lower sensitivity to certain beta-lactams (Hartman & Tomasz, 1984; Wielders et al., 2002). Similarly, a typically non-essential *Enterococcus faecalis* bPBP with low reactivity to cephalosporins provides functional redundancy to the primary division-associated bPBP during cephalosporin treatment (Djorić et al., 2020; Nelson et al., 2024).

While the molecular mechanisms enabling redundancies between bPBPs remain unclear, these observations suggest a few possibilities. In the case of *S. enterica*, the two division-specific bPBPs appear to function independently since neither bPBP requires the presence of the other to function in their respective niche (Castanheira et al., 2017). Similarly, elongasome function in *B. subtilis* can be supported by the activity of one of two distinct bPBPs, PBP2a*_Bs_* and PBPH*_Bs_* (Wei et al., 2003). Notably, elongasome dynamics are unaffected in cells lacking either of the two bPBPs (Middlemiss et al., 2024), so both bPBPs presumably can complex with the elongation-specific SEDS protein and regulate its function. In contrast, the functional redundancy reported between the division-associated bPBPs, PBP3*_Bs_* and PBP2b*_Bs_* requires the presence of the catalytically inactivated PBP2b*_Bs_* protein (Sassine et al., 2017). Since SEDS-bPBP pairs are generally thought to be comprised of cognate partners whose activities are coordinated by and require their interaction (Shlosman et al., 2023; Sjodt et al., 2020; Taguchi et al., 2019), the requirement of PBP2b*_Bs_* is likely dictated by the structural role it plays in interacting with and stimulating the glycosyltransferase activity of the division-specific SEDS protein, FtsW. This is consistent with an *in vitro* study, which showed that a catalytically inactive form of a bPBP can support PG polymerization by its SEDS partner (Taguchi et al., 2019). Furthermore, the redundancy observed in *E. faecalis* is suggested to require direct interaction between the two bPBPs within the larger divisome complex (Nelson et al., 2024). In contrast, since the two division-specific bPBPs in *S. enterica* can function independently from one another, it is possible that either enzyme can interact with the division-specific SEDS glycosyltransferase, FtsW (Castanheira et al., 2017).

Since the activity of *C. difficile* PBP3 during asymmetric division requires the presence of SpoVD (**Figure 1**), the redundancy between the sporulation-specific SpoVD and PBP3 bPBPs is more similar to the catalytic redundancy observed between division-specific bPBPs in *B. subtilis* likely due to SpoVD allosterically activating SpoVE’s activity. SpoVD may also promote the formation and function of the polar divisome complex because PBP3 interacts with SpoVD and potentially additional members of the polar divisome during sporulation (**Figure 5**). The active recruitment of functionally redundant bPBPs to the divisome is consistent with the finding that septal PG synthesis is a highly controlled process involving various localization and regulatory mechanisms (Egan et al., 2020). Indeed, due to the requirement for SpoVD, we speculate that PBP3 is likely recruited as an additional factor rather than replacing SpoVD in the polar divisome complex. While confirmation and characterization of the mechanisms and factors facilitating this recruitment require further study, our results suggest that PBP1’s recruitment to the polar divisome may be important for helping *spoVD* catalytic mutant strains complete asymmetric division. This hypothesis is based on our observations that (i) PBP3’s catalytic activity is dispensable for promoting asymmetric division in a *spoVD* catalytic mutant strain (**Figure 4**) and (ii) PBP1 appears to interact directly with PBP3 in two-hybrid analyses and co-immunoprecipitates with both SpoVD and PBP3 purified from sporulating *C. difficile* cells (**Figure 5**).

It remains unclear why SpoVD catalytic activity is largely dispensable during asymmetric division but essential for cortex synthesis. It is possible that cortex synthesis involves SpoVE-SpoVD to function as part of a larger PG synthetic complex, similar to the divisome and elongasome. Since SpoVE-SpoVD function during this process requires their specific localization to the forespore, it is likely that other factors are involved in their regulation. It is, therefore, possible that the inability of PBP3 to functionally compensate for the loss of SpoVD catalytic activity during cortex synthesis results from its decreased affinity for these unknown factors. Another possibility is that the amount of crosslinking required is higher than the amount PBP3 is able to supplement during this process.

Finally, redundancy in bPBP activity is likely widespread among spore-forming bacteria based on our finding that they typically encode multiple additional bPBPs, the number of which often exceeds the number of SEDS glycosyltransferases encoded in the genome (**Figure 6**).

Defining specialized roles or redundancies between these enzymes requires further study and may have important implications for the ability of these organisms to respond to environmental and antibiotic stress. Interestingly, while most *C. difficile* strains carry five distinct HMW PBPs, a study characterizing clinical strains identified an additional bPBP encoded by *Clostridium difficile* ribotype 017 isolates (Isidro et al., 2017, 2018). Some of these isolates have a lower sensitivity to the beta-lactam imipenem and carry mutations in the transpeptidase catalytic sites of the division-associated PBP1 and elongation-associated PBP2. The additional bPBP encoded by these strains, therefore, may additionally contribute to imipenem resistance by providing functional redundancy to these enzymes or SpoVD, recapitulating the scenario observed in MRSA strains. Taken together with our findings, these observations highlight the need to define functional redundancies between PBPs in *C. difficile* and their possible roles in driving antibiotic resistance of this important pathogen.

## Methods

### C. difficile strain construction and growth conditions

All *C. difficile* strains are derived from the 630Δ*erm* strain. Deletion and complementation strains were constructed in a Δ*pyrE* background strain using *pyrE*-based allele-coupled exchange as previously described (Ng et al., 2013). All strains used in the study are reported in Table S1.

Strains were grown at 37°C under anaerobic conditions using a gas mixture containing 85% N_2_, 5% CO_2_, and 10% H_2_.

### E. coli strain constructions

Table S2 lists all plasmids used in the study, with links to plasmid maps containing all primer sequences used for cloning. Plasmids were cloned via Gibson assembly, and cloned plasmids were transformed into *E. coli* (DH5α or XL1-Blue strains). All plasmids were confirmed by sequencing the inserted region. Confirmed plasmids were transformed into the *E. coli* HB101(pRK24) strain for conjugation with *C. difficile* when needed. All *E. coli* HB101 strains used for conjugation are indicated listed in Table S2.

### Plate-based sporulation assays

For assays requiring sporulating cells, cultures were grown to early stationary phase, back-diluted >25-fold into BHIS, and grown until they reached exponential phase (OD_600_ between 0.35 and 0.75). 120 µL of exponentially growing cells were spread onto 70:30 (70% SMC media and 30% BHIS media) agar plates (40 mL media per plate). After 18-22 hours of growth, sporulating cells were collected into phosphate-buffered saline (PBS), and sporulation levels were visualized by phase-contrast microscopy as previously described (Pishdadian et al., 2015).

### Heat resistance assay

Heat-resistant spore formation was measured 20-22 hours after sporulation induction on 70:30 agar plates by resuspending sporulating cells in PBS, dividing the sample into two, heat-treating one of the samples at 60°C for ∼30 min, and comparing the colony-forming units (CFUs) in the untreated sample to the heat-treated sample (Fimlaid et al., 2015). Heat-resistance efficiencies represent the average ratio of heat-resistant CFUs to total CFUs for a given strain relative to the average ratio for the wild-type strain.

### Fluorescence and phase-contrast microscopy

Fluorescence microscopy was performed on sporulating cells using Hoechst 33342 (Molecular Probes; 15 µg/mL) and FM4-64 (Invitrogen; 1 µg/mL) to stain nucleoid and membrane, respectively. All samples for a given experiment were imaged from a single agar pad (1.5% low-melting point agarose in PBS).

Phase-contrast images in **Figure 3** were obtained using a Zeiss Axioskop upright microscope with a 100× Plan-NEOFLUAR oil-immersion phase-contrast objective and a Hamamatsu C4742-95 Orca 100 CCD Camera. All other phase-contrast and fluorescence images were acquired using a Leica DMi8 inverted microscope with a 63× 1.4 NA Plan Apochromat oil-immersion phase-contrast objective, a high precision motorized stage (Pecon), and an incubator (Pecon) set at 37°C. Excitation light was generated by a Lumencor Spectra-X multi-LED light source with integrated excitation filters. An XLED-QP quadruple-band dichroic beam-splitter (Leica) was used (transmission: 415, 470, 570, and 660 nm) with an external filter wheel for all fluorescent channels. FM4-464 was excited at 550/38 nm and emitted light was filtered using a 705/72 nm emission filter (Leica); Hoechst was excited at 395/40 nm and emitted light was filtered using a 440/40 nm emission filter (Leica). Emitted and transmitted light was detected using a Leica DFC 9000 GTC sCMOS camera. 1 to 2 µm z-stacks were taken when needed with 0.21 µm z-slices.

Images were acquired and exported using the LASX software without further processing. After export, images were processed using Fiji (Schindelin et al., 2012) to remove out-of-focus regions, and the best-focused z-planes for all channels were manually selected. Image scaling was adjusted to improve brightness and contrast for display and was applied equally to all images shown in a single panel. Visualization of quantified data and any associated statistical tests were performed using Prism 10 (GraphPad Software, San Diego, CA, USA).

### Transmission electron microscopy

Sporulating cells were collected ∼22 h after sporulation induction on 70:30 or SMC agar plates. Cells were fixed and sent for processing for electron microscopy by the University of Vermont Microscopy Center, as previously described (Putnam et al., 2013; Shrestha et al., 2023). All TEM images were captured on a JEOL 1400 Transmission Electron Microscope (Jeol USA, Inc., Peabody, MA) with an AMT XR611 high-resolution 11-megapixel mid-mount CCD camera.

### Western blot analysis

Samples were collected 18-22 hours after sporulation induction on 70:30 agar plates and processed for immunoblotting. Sample processing involved multiple freeze-thaws in PBS followed by the addition of EBB buffer (9 M urea, 2 M thiourea, 4% SDS, 2 mM b- mercaptoethanol), boiling, pelleting, resuspension, and boiling again before loading on a gel. All proteins were resolved using 4–15% precast polyacrylamide gels (Bio-Rad) and transferred to polyvinylidene difluoride membranes, which were subsequently probed with rabbit (anti-PBP3 (this study) and anti-SpoVD (Shrestha et al., 2023); both at 1:1,000 dilution), mouse (anti-Spo0A (Fimlaid et al., 2013) at 1:1,000 dilution) and chicken (anti-GDH at 1:5,000 dilution) polyclonal primary antibodies, and anti-rabbit (IR800 or IR680), anti-mouse (IR680) and anti-chicken (IR800) secondary antibodies (LI-COR Biosciences, 1:20,000 dilution). Blots were imaged using a LiCor Odyssey CLx imaging system. The results shown are representative of multiple experiments.

### Antibody production

Anti-PBP3 and anti-PBP1 antibodies used for western blots in this study were raised in rabbits by Cocalico Biologicals against PBP3 and PBP1 variants lacking the transmembrane domains. PBP3_Δ1-35_-His_6_ and His_6_-PBP1_Δ1-77_ were produced in BL21(DE3) *E. coli* harboring pET28a-*pbp3* _Δ1-35_-His_6_ and pET28a-His_6_-*pbp1*_Δ1-77_, respectively, and purified by Ni^2+^-affinity purification as previously described (Fimlaid et al., 2013). Antisera reactivity and specificity for PBP3 and PBP1 were validated by western blot against *C. difficile* lysate from WT and either a Δ*pbp3* mutant (this work) or a *pbp1* CRISPR-interference knock-down strain (Shrestha et al., 2023), respectively.

### Bacterial two-hybrid analyses

Bacterial adenylate cyclase two-hybrid (BACTH) assays were conducted as previously described (Karimova et al., 1998) using *E. coli* BTH101 cells. Briefly, BTH101 cells were transformed with 100 ng of each plasmid and plated on LB agar plates supplemented with 50 µg/ml kanamycin, 100 µg/ml Ampicillin, and 0.5 mM isopropyl β-D-thiogalactopyranoside (IPTG).

Plates were incubated for 64-68 hours at 30°C, and β-galactosidase activity was quantified in Miller units as previously detailed (Dahlstrom et al., 2015). The β-galactosidase activity of cells transformed with the empty pUT18C and pKT25 vectors was used as a negative control for normalization.

### Co-immunoprecipitation (Co-IP) analyses

To immunoprecipitate FLAG-tagged proteins from sporulating *C. difficile*, exponentially growing cultures of *C. difficile* harboring pbp3-FLAG3 or spoVD-FLAG3 expression constructs and appropriate control strains were spread onto 70:30 agar plates. After 12 hours of growth on 70:30 plates, sporulating cells were scraped from 3 plates per strain and pooled in 1 mL of PBS. A portion of cells was visualized by microscopy with peptidoglycan labeling to confirm the presence of sporulating cells undergoing asymmetric division in each sample. Cells were crosslinked with 0.25% final concentration of PFA for 15 minutes at 37°C and quenched with 350 mM glycine for 10 minutes on ice. Crosslinked cells were then pelleted, re-suspended in 750 µL of FLAG IP Buffer (150 mM NaCl, 50 mM Tris-HCl pH 7.5), transferred to screwcap tubes containing MP Biomedicals Lysing Matrix E, and frozen at -80°C. Frozen cells were thawed and bead beat on an MP Biomedicals FastPrep-24 four times at 5.5 M/second for 1 minute, resting on ice for 5 minutes between rounds of bead-beating. Next, 1X HALT protease inhibitors and dodecyl-β-d-maltoside (DDM) detergent were added to lysed cells to a final concentration of 0.5%, and mixtures were rotated at room temperature for 1 hour to solubilize membrane proteins. Lysates were clarified by centrifuging at 10,000 × g for 1 minute, and 200 µL of pre-equilibrated Anti-FLAG M2 Magnetic Dynabead resin was added to the clarified lysate. The lysate-resin mixture was rotated at room temperature for 1 hour. To remove unbound proteins, the resin was washed three times briefly and once for 5 minutes with 1 mL FLAG IP Buffer containing 0.5% DDM. Then, the resin was washed three times briefly and once for 5 minutes with FLAG IP Buffer containing no detergent. This step was repeated a total of two times. Finally, bound proteins were eluted from the resin with 2 μg/mL 3XFLAG peptide, boiled in 1X sample buffer for 10 minutes to reverse crosslinks, and then analyzed by western blot using rabbit polyclonal antibodies against PBP1 (TF134, this study), PBP3 (TF135, this study), SpoVD (TF124, (Shrestha et al., 2023)), SpoIID (TF121, (Ribis et al., 2018)), or a mouse monoclonal M2 anti-FLAG antibody (Sigma).

## Acknowledgments

We acknowledge the University of Vermont Microscopy Imaging Core for processing the samples and acquiring the images for all Transmission Electron Microscopy analyses. We are grateful to members of the Shen lab for helpful discussions and feedback on the project. The National Institute of Allergy and Infectious Diseases grant R01 AI122232 (to A.S.) and Burroughs Wellcome Fund for Investigators in Pathogenesis Award (to A.S.) provided funding for this work. Additionally, G.A.H. was supported by the Tufts University Institutional Research Career and Academic Development Award Program funded by the National Institute of General Medical Sciences K12 GM133314.

## Author Contributions

S.S. and A.S. conceived the study. S.S., J.M.D., G.A.H., M.M, and A.S. performed and analyzed experiments. A.S. supervised the study. S.S. and A.S. wrote the manuscript with input from the other authors. All authors reviewed and approved the manuscript.

