## Supplementary figures and tables for "Functional redundancy between penicillin-binding proteins during asymmetric cell division in *Clostridioides difficile*"

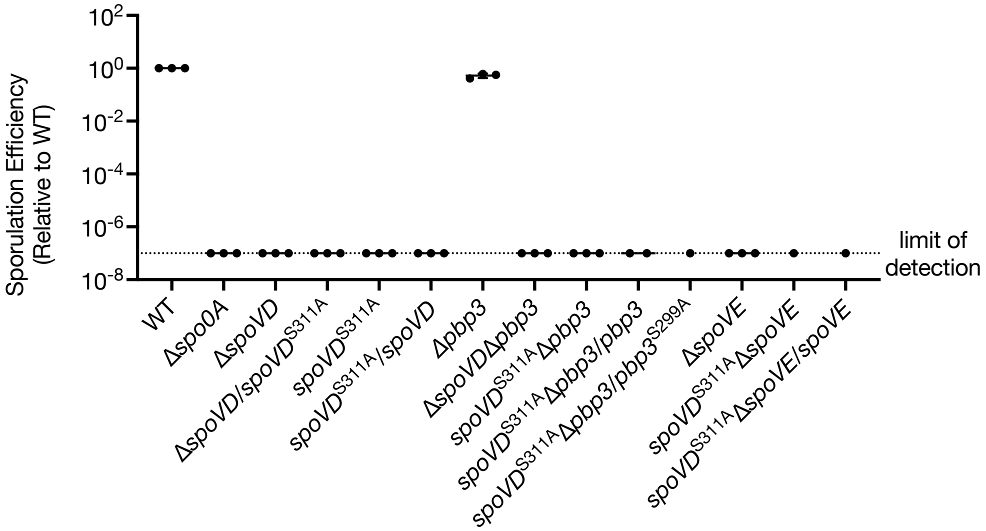


**Fig. S1 | The efficiency of heat-resistant spore formation (sporulation efficiency) of various mutant and complemented strains relative to WT.** Means with standard deviation are indicated. Cells were collected from sporulation-inducing 70:30 plates ~20-22 hours after inoculation.

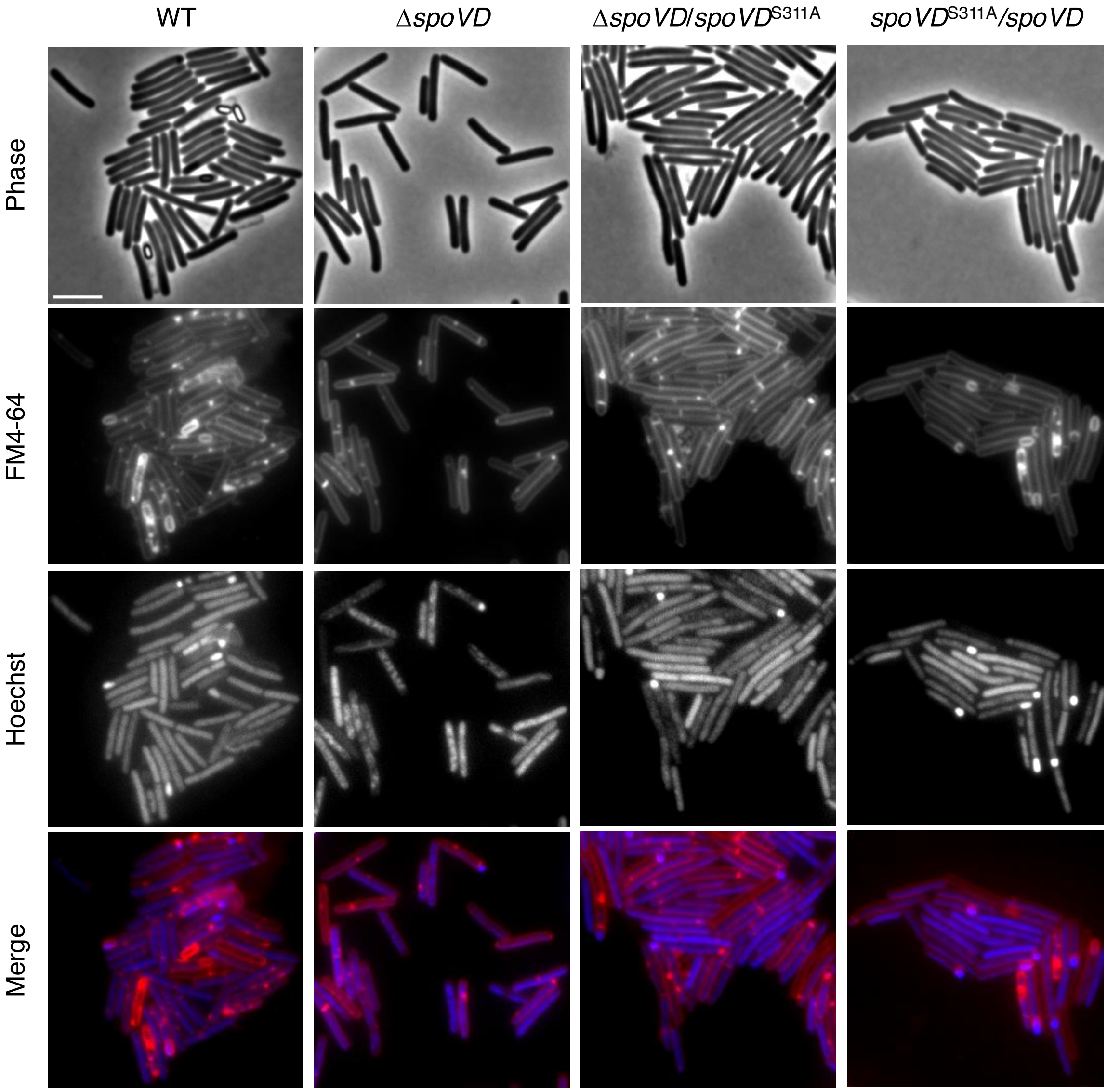


**Figure S2. Cytological profiling of strains analyzed in Figure 1**. Representative phase-contrast and fluorescence micrographs of the indicated strains sampled after ~20 hours of growth on sporulation-inducing 70:30 plates. The nucleoid was stained using Hoechst, and the cell membrane was stained using FM4-64. Scale bar = 5 µm.

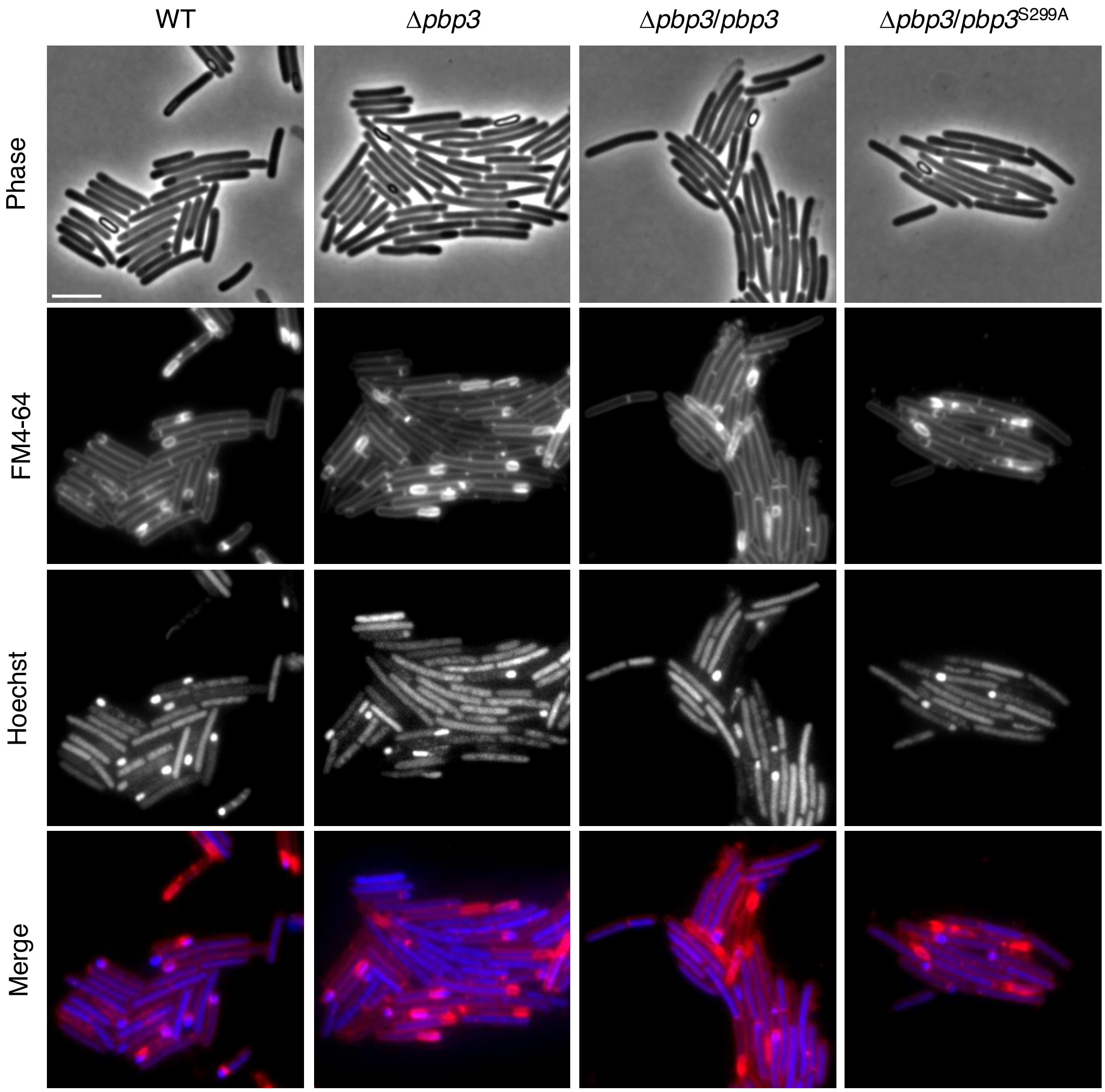


**Figure S3. Cytological profiling of strains analyzed in Figure 3**. Representative phase-contrast and fluorescence micrographs of the indicated strains sampled after ~20 hours of growth on sporulation-inducing 70:30 plates. The nucleoid was stained using Hoechst, and the cell membrane was stained using FM4-64. Scale bar = 5 µm.

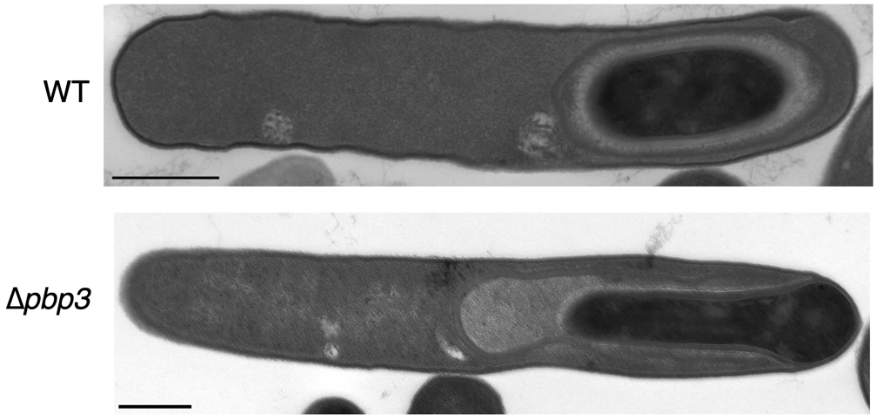


**Figure S4. Transmission Electron Microscopy of Δ*pbp3* cells.** Representative transmission electron micrographs (TEM) of WT and Δ*pbp3* strains. The forespore of Δ*pbp3* cells was frequently observed to form an elongated shape in the TEM samples. Scale bar = 500 nm.

**Table S1.** ***C. difficile*** **strains used in this study**

| Strain number | Strain description | Description | Study |
|---|---|---|---|
| 756 | 630Δ*erm*Δ*pyrE* | *erm*-sensitive derivate of 630 with a deletion in *pyrE* | Ng *et al*. 2013 |
| 846 | 630Δ*erm*-p | 630Δ*erm* with *pyrE* restored in the native locus - used as the WT strain in the study | Donnelly *et al*. 2017 |
| 849 | 630Δ*erm*-p Δ*spo0A* | 630Δ*erm* with a deletion of *spo0A* and *pyrE* restored | Donnelly *et al*. 2017 |
| 2595 | 630Δ*erm*-p Δ*spoVD* | 630Δ*erm* with a deletion of *spoVD* (*cd630_26560*) and *pyrE* restored | Shrestha *et al*. 2023 |
| 2734 | 630Δ*erm*-p Δ*pbp3*Δ*pyrE* | 630Δ*erm*Δ*pyrE* with a deletion of *pbp3* (*cd630_12290*) | Shrestha *et al*. 2023 |
| 2948 | 630Δ*erm*-p Δ*pbp3* | 630Δ*erm*Δ*pbp3* with *pyrE* restored | Shrestha *et al*. 2023 |
| 3004 | 630Δ*erm*-p Δ*spoVD/spoVD* | 630Δ*erm*Δ*spoVD* with *pyrE* restored and *spoVD* complemented in the *pyrE* locus | Shrestha *et al*. 2023 |
| 3022 | 630Δ*erm*-p Δ*pbp3/pbp3* | 630Δ*erm*Δ*pbp3* with *pyrE* restored and *pbp3* complemented in the *pyrE* locus | This study |
| 3031 | 630Δ*erm*-p Δ*spoVE* | 630Δ*erm*Δ*spoVE* with *pyrE* restored | Shrestha *et al*. 2023 |
| 3072 | 630Δ*erm*-p Δ*spoVE/spoVE* | 630Δ*erm*Δ*spoVE* with *pyrE* restored and *spoVE* complemented in the *pyrE* locus | Shrestha *et al*. 2023 |
| 3326 | 630Δ*erm*Δ*pyrE*Δ*spoVD*Δ*pbp3* | 630Δ*erm*Δ*pyrE* with a sequential deletion of *spoVD* and *pbp3* | This study |
| 3581 | 630Δ*erm*-p Δ*spoVD*Δ*pbp3* | 630Δ*erm*Δ*spoVD*Δ*pbp3* with *pyrE* restored | This study |
| 4110 | 630Δ*erm*Δ*pyrE spoVD*$^{S311A}$ | 630Δ*erm*Δ*pyrE* with the native *spoVD* mutated (serine to alanine substitution (tct>gct) in residue 311) | This study |
| 4348 | 630Δ*erm*-p *spoVD*$^{S311A}$ | 630Δ*erm*Δ*pyrE spoVD*$^{S311A}$ with *pyrE* restored | This study |
| 4351 | 630Δ*erm*-p *spoVD*$^{S311A}$/*spoVD* | 630Δ*erm*Δ*pyrE spoVD*$^{S311A}$ with *pyrE* restored and *spoVD* complemented in the *pyrE* locus | This study |
| 4361 | 630Δ*erm*-p *spoVD*$^{S311A}$ Δ*pbp3* | 630Δ*erm*Δ*pyrE spoVD*$^{S311A}$ with deletion of *pbp3* and *pyrE* restored | This study |
| 4390 | 630Δ*erm*-p *spoVD*$^{S311A}$ Δ*pbp3/pbp3* | 630Δ*erm*Δ*pyrE spoVD*$^{S311A}$Δ*pbp3* with *pyrE* restored and *pbp3* complemented in the *pyrE* locus | This study |
| 4432 | 630Δ*erm*-p Δ*pbp3/pbp3*$^{S299A}$ | 630Δ*erm*Δ*pyrE* Δ*pbp3* with *pyrE* restored and mutated *pbp3* (serine to alanine substitution (tct>gct) in residue 299) in the *pyrE* locus | This study |
| 4441 | 630Δ*erm*-p Δ*pbp3/pbp3*$_{3XFLAG}$ | 630Δ*erm*Δ*pyrE* Δ*pbp3* with *pyrE* restored and *pbp3-3XFLAG* in *pyrE* locus | This study |
| 4568 | 630Δ*erm*-p Δ*spoVD/spoVD*-$_{3XFLAG}$ | 630Δ*erm*Δ*pyrE* Δ*spoVD* with *pyrE* restored and *spoVD-3XFLAG* in *pyrE* locus | This study |
| 4574 | 630Δ*erm*-p *spoVD*$^{S311A}$ Δ*pbp3/pbp3*$^{S299A}$ | 630Δ*erm*Δ*pyrE spoVD*$^{S311A}$Δ*pbp3* with *pyrE* restored and *pbp3*$^{S299A}$ complemented in the *pyrE* locus | This study |
| 4623 | 630Δ*erm*-p *spoVD*$^{S311A}$ Δ*spoVE* | 630Δ*erm*Δ*pyrE spoVD*$^{S311A}$ with deletion of *spoVE* and *pyrE* restored | This study |
| 4626 | 630Δ*erm*-p *spoVD*$^{S311A}$ Δ*spoVE/spoVE* | 630Δ*erm*Δ*pyrE spoVD*$^{S311A}$Δ*spoVE* with *pyrE* restored and *spoVE* complemented in the *pyrE* locus | This study |

**Table S2.** ***E. coli*** **strains used in this study**

| Strain number | Strain description | Relevant genotype or link to annotated plasmid sequence | Source |
|---|---|---|---|
| 41 | DH5α | $F^-$ Φ80*lacZ*ΔM15 Δ(*lacZYA-argF*) U169 *recA1 endA1 hsdR17* ($rK^-$, $mK^+$) *phoA supE44* λ– *thi-1 gyrA96 relA1* | D. Cameron |
| 531 | HB101 | $F^-$ *mcrB mrr hsdS20*($rB^-mB^-$) *recA13 leuB6 ara-13 proA2 lavYI galK2 xyl-6 mtl-1 rpsL20* carrying pRK24 | C. Ellermeier |
| 1218 | BTH101 | $F^-$, *cya-99, araD139, galE15, galK16, rpsL1* ($Str^r$), *hsdR2, mcrA1, mcrB1* | Euromedex |
| 1452 | XL1-Blue | *recA1 endA1 gyrA96 thi-1 hsdR17 supE44 relA1 lac* carrying F *proAB lacIqZΔM15 Tn10* ($Tet^r$) | C. Huston |
| 1539 | DH5α / pMTL-YN3 | pMTL-YN3 in DH5α | Ng *et al.* 2013 |
| 1662 | HB101 / pMTL-YN1C | pMTL-YN1C in HB101/pRK24 | Ng *et al.* 2013 |
| 2469 | HB101 pMTL-YN3 Δ*pbp3* | https://benchling.com/s/seq-ZXq5K09mLbsPHse4YNTa | Shrestha *et al.* 2023 |
| 2475 | HB101 pMTL-YN1C-*spoVD* | https://benchling.com/s/seq-rB133tLjJlc7rJJXQtZs | Shrestha *et al.* 2023 |
| 2627 | HB101 pMTL-YN1C-*spoVD*$^{S311A}$ | https://benchling.com/s/seq-RlSZsqjrdgU9pUnI101G | This study |
| 2631 | HB101 pMTL-YN1C-*pbp3* | https://benchling.com/s/seq-Y7cfuxWOsm45Pz7Q8veh?m=slm-OWqt0kN1tH15a8vtNByM | This study |
| 2672 | HB101 pMTL-YN1C-*spoVE* | https://benchling.com/s/seq-J2BDth9Xje1Ry0RYL8yY | Shrestha *et al.* 2023 |
| 3354 | HB101 pMTL-YN3-*spoVD*$^{S311A}$ | https://benchling.com/s/seq-RlSZsqjrdgU9pUnI101G?m=slm-igkxB8eqZvrVXk97wRLj | This study |
| 3568 | BL21(DE3) pET28a-*pbp1*$_{\Delta 1\text{-}78}$ | https://benchling.com/s/seq-H4Q8cuaGUBznJVmcUxWr?m=slm-0jWzqAC1NRqpR0XxXAPV | This study |
| 3611 | HB101 pMTL-YN1C-*spoVD*$^{3XFLAG}$ | https://benchling.com/s/seq-8ozjvbJl1aFC4yvUZ93R?m=slm-H4bxr5oIsG57MvbITzcV | This study |
| 3697 | HB101 pMTL-YN1C *pbp3*$^{S299A}$ | https://benchling.com/s/seq-W0iG7JIhBgkWkIY30JSn?m=slm-Da99hzZ1nSKhBu3nl8c6 | This study |
| 3699 | HB101 pMTL-YN1C *pbp3*$^{3XFLAG}$ | https://benchling.com/s/seq-84uLyNFxTAf0F0dmy7W7?m=slm-PI4ps1rk6G0Rmkce22Rx | This study |
| 3709 | BL21(DE3) pET28a *pbp3*$_{\Delta TM}$-His$_6$ | https://benchling.com/s/seq-Na7IhwYzfBMC3cgvWEDq?m=slm-flacnqoNYZnT4Bn4mRLN | This study |
| 3151 | XL1-Blue pUT18C-*spoVD* | https://benchling.com/s/seq-CoDkpnu9UjIw1NGMf7K3?m=slm-c2x8L4sgsvccomDfHd95 | Shrestha *et al.* 2023 |
| 3152 | XL1-Blue pKT25-*spoVD* | https://benchling.com/s/seq-WE1eUEvUa9oPe1SHOONt?m=slm-ehUZnAOhmYE9ik8Lc6oJ | Shrestha *et al.* |
| 3153 | XL1-Blue pUT18C-*spoVE* | https://benchling.com/s/seq-J6xY0qAz4gLUCvIsvPtr?m=slm-Vel4vhorTutATD2xLKCg | Shrestha *et al.* 2023 |
| 3154 | XL1-Blue pKT25-*spoVE* | https://benchling.com/s/seq-q0Hi3dYXXUh38AIJmwLo?m=slm-PfOO6WHj8YBpHHf4cZNm | Shrestha *et al.* 2023 |
| 3155 | XL1-Blue pUT18C-*ftsL* | https://benchling.com/s/seq-bmQB9ktQ33MqNQkBLm55?m=slm-tVTkjwNmnkSaVBZQJ9df | Shrestha *et al.* 2023 |
| 3156 | XL1-Blue pKT25-*ftsL* | https://benchling.com/s/seq-EMRCycRB9UdqQCbeX9jT?m=slm-Eu8SRXD4GMNsWRo31JCl | Shrestha *et al.* 2023 |

| | | | |
|---|---|---|---|
| 3157 | XL1-Blue pUT18C-*ftsQ* | https://benchling.com/s/seq-iL0HyX5G3dtQADBqOFfK?m=slm-ukilYA8jsYlCh4y62YLT | Shrestha *et al.* |
| 3158 | XL1-Blue pKT25-*ftsQ* | https://benchling.com/s/seq-JEX75QVog9sFurh310Wa?m=slm-jc3ZO90jWPfMC6Hgadz6 | Shrestha *et al.* |
| 3159 | XL1-Blue pUT18C-*ftsB* | https://benchling.com/s/seq-WvgT8dJSLtfQwy4WClZt?m=slm-iwF6SYpgknb2XL5Iaw1h | Shrestha *et al.* |
| 3160 | XL1-Blue pKT25-*ftsB* | https://benchling.com/s/seq-aV3xwMOTrLh41216a6WR?m=slm-xGsUs6j0EwZZPQjGb5eB | Shrestha *et al.* |
| 3161 | XL1-Blue pUT18C-*pbp3* | https://benchling.com/s/seq-PAZ0xi4hBP1lUJVGzN35?m=slm-bv2a9iCl43AKUPloC1fi | This study |
| 3162 | XL1-Blue pKT25-*pbp3* | https://benchling.com/s/seq-FXTvwywfrSVha33A3rQe?m=slm-5bR9Y51YRgxObZ2L9PjY | This study |
| 3339 | XL1-Blue pUT18C-*pbp1* | https://benchling.com/s/seq-XfC81UyoBHmvnPTxis0X?m=slm-7CejXGO6LHOxZbGA8qdy | This study |
| 3340 | XL1-Blue pKT25-*pbp1* | https://benchling.com/s/seq-iMQhOTXvze4hhDbDFemu?m=slm-2Fv0U9MQL2rEljCw7Vtx | This study |
| 3341 | XL1-Blue pUT18C-*pbp2* | https://benchling.com/s/seq-VHP42dY1B34apZNcX1Y2?m=slm-m6OusVY9cLFCDRsfhlZG | This study |
| 3342 | XL1-Blue pKT25-*pbp2* | https://benchling.com/s/seq-0bkQGwHfBPiVSQ50x9ap?m=slm-9N3vZHIFmrFuKyKcKoKl | This study |
| 3343 | XL1-Blue pUT18C-*rodA* | https://benchling.com/s/seq-zmCqlG5VjhoGcCEaWqDf?m=slm-6zDIR9ELx56d1AZJqLvD | This study |
| 3344 | XL1-Blue pKT25-*rodA* | https://benchling.com/s/seq-bIZxKmimHnx1eDrmKMvJ?m=slm-GyIHHOHSX9wgdFMbhSsk | This study |
